## Supplemental Text and Figures for "Genomic determinants of soil depth niche partitioning in Gagatemarchaeaceae, a novel family of deeply-rooted Thaumarchaeota"

#### Classifications

Candidatus "*Gagatemarchaeum stordalenia*" (sp. nov., gen. nov). "Gagatem" refers to its prevalence in peat environments. The name "stordalenia" originates from the geographical origin (Stordalen Mire, Sweden) of the reference (type) genome, bog-1369. It encodes genes for aerobic respiration and likely uses organic substrates, such as carbohydrates, peptides and fatty acids for organoheterotrophic growth. It is currently not cultured and known only from environmental sequencing. Genomes of this genera possess a high GC-content (around 60%) and genome size of its members range from 2.2-3.4 Mb.

Candidatus "*Subgagatemarchaeum marcellia*" (sp. nov., gen. nov). "Subgagatem" refers to its prevalence in subsoils and peat environments. The name "'marcellia" originates from the geographical origin (Marcell Experimental Forest, MN, USA) of the reference (type) genome, Fn1. It likely uses organic substrates, such as carbohydrates, peptides and fatty acids for organoheterotrophic growth. While the type genome, Fn1, encodes the genes for microaerophilic respiration cytochrome bd ubiquinol oxidase, the majority of studied genomes from this genera encode the aerobic respiration heme-copper oxygen reductases. It is currently not cultured and only known from environmental sequencing. Genomes of this genera possess a high GC-content (around 58%) and genome size of its members range form 1.2-2.5 Mb.

Description of Gagatemarchaeaceae (fam. nov). This family was previously referred as the Group I.1c Thaumarchaeota. Description is the same as for the genus *Gagatemarchaeum*. Suff. -aceae, ending to denote family. Type genus *Gagatemarchaeum* gen. nov.

### Supplementary methods

#### Gene marker selection and phylogenomic inference

For each dataset, ortholog groups (OGs) were detected using Roary (-i 50, -iv 1.5)[1]. Core OGs were defined as those present in single copy in each genome and present in at least 70% of the genomes. Core OGs were aligned individually using MAFFT L-INS-i[2], and spurious sequences and poorly aligned regions were removed with trimAl (automated1, resoverlap 0.55 and seqoverlap 60)[3]. Alignments were removed from further analysis if they presented evidence of recombination using the PHItest[4]. The remaining alignments were concatenated into a supermatrix for each dataset.

Maximum-likelihood trees were constructed for each dataset supermatrix with IQ-TREE 2.0.3[5] using the complex mixture model LG+C60+G+F. Branch supports were computed using the SH-aLRT test[6] and 2,000 UFBoot replicates. A hill-climbing nearest-neighbour interchange (NNI) search was performed to reduce the risk of overestimating branch supports.

#### Functional annotation of genomes

Genomes were annotated with the KEGG database[7] using GhostKOALA[8], with the arCOG database [9] using Diamond BLASTp[10] (best-hit and removing matches with e-value >10−5, % ID < 35, alignment length <80 or bit score <100) and with the Pfam[11] database using hmmsearch[12] (-T 80). The subfamily classification of *cyd*A was performed using hmmsearch (-T 80) with the *cyd*A subfamily database[13]. The subfamily classification of *cox*A genes was performed using the heme-copper oxygen reductase database[14]. Carbohydrate-active enzymes were annotated using profile HMM from dbCAN (http://bcb.unl.edu/dbCAN2/) (filtered with hmmscan-parser.sh and by removing matches with mean posterior probability <0.7). Peptidases were annotated using Pfam profile HMMs corresponding to MEROPs families, as described previously[15]. Extracellular carbohydrate-active enzymes peptidases were identified using Signalp 5.0[16] (-org arch, archaeal signal peptides) to detect the presence of signal peptides.

The presence of motility genes in Gagatemarchaeaceae was initially assessed by the presence of the conserved archaellum subunits C (arCOG05119), D/E (arCOG02964), F (arCOG01824), G (arCOG01822) and J (arCOG01809). However, previous work on Poseidoniales indicated that some archaea might possess divergent motility loci[15]. Therefore, the profile HMM of archaeal flagellin (PF01917) gene was used as markers for possible divergent motility, even when other archaellum related genes were absent.

The 5S, 16S and 23S rRNA and tRNA genes were identified using Barrnap v0.9 (--kingdom arc, archaeal rRNA) (https://github.com/tseemann/barrnap) and tRNAscan-SE v2.0.5[17] (-A, archaeal tRNA), respectively. The 16S rRNA genes from the different genomes were compared by a pairwise analysis using BLASTn v2.9.0[18].

#### Supplementary Figures

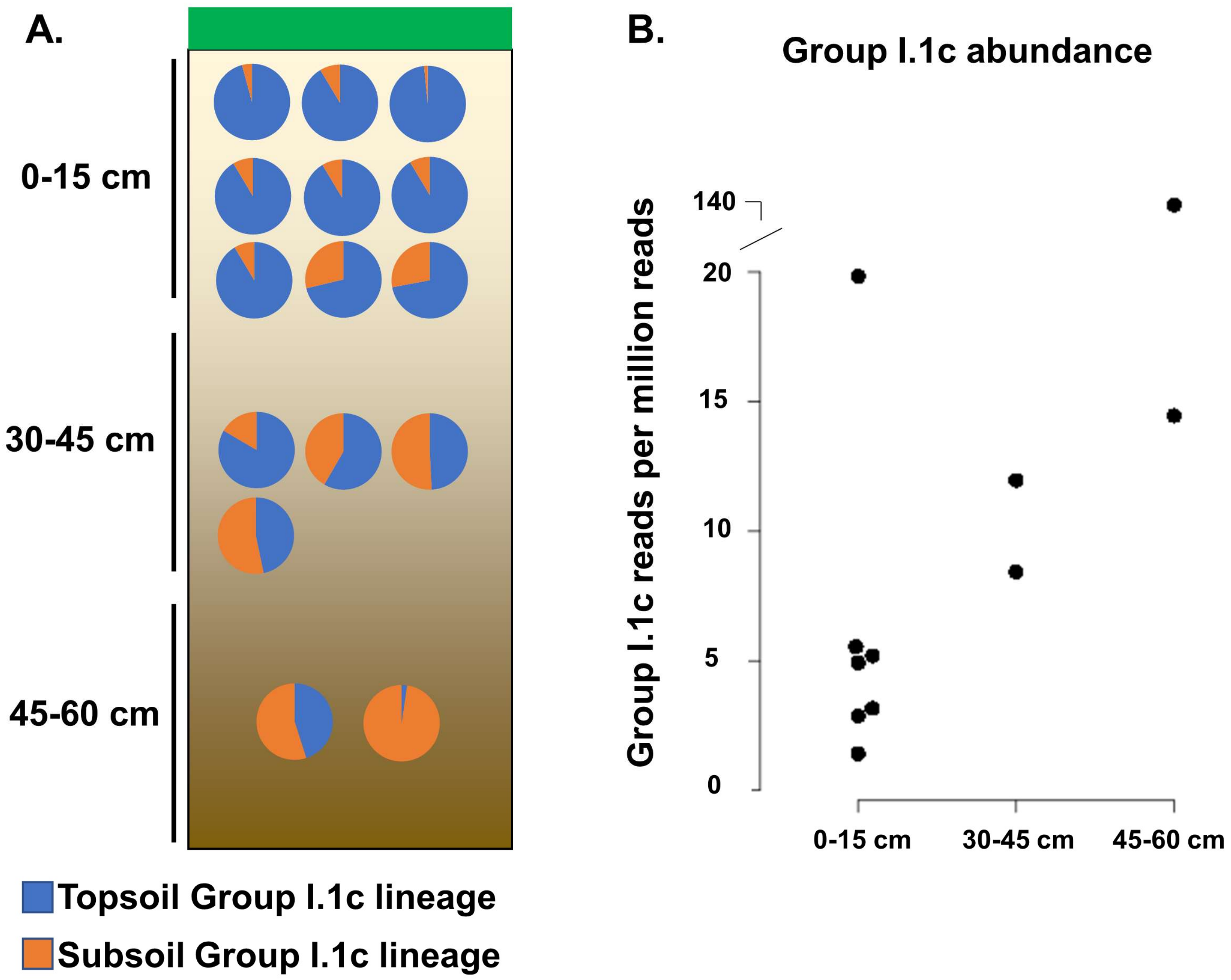


**Supplementary Fig. 1. Depth distribution of soil Group I.1c lineages.** (A) The Group I.1c community composition changes from Topsoil lineage dominated to Subsoil lineage dominated with increasing depth, and (B) the overall abundance of Group I.1c increases with depth. Metagenomic reads were recruited to genomes of the Topsoil and Subsoil lineages.

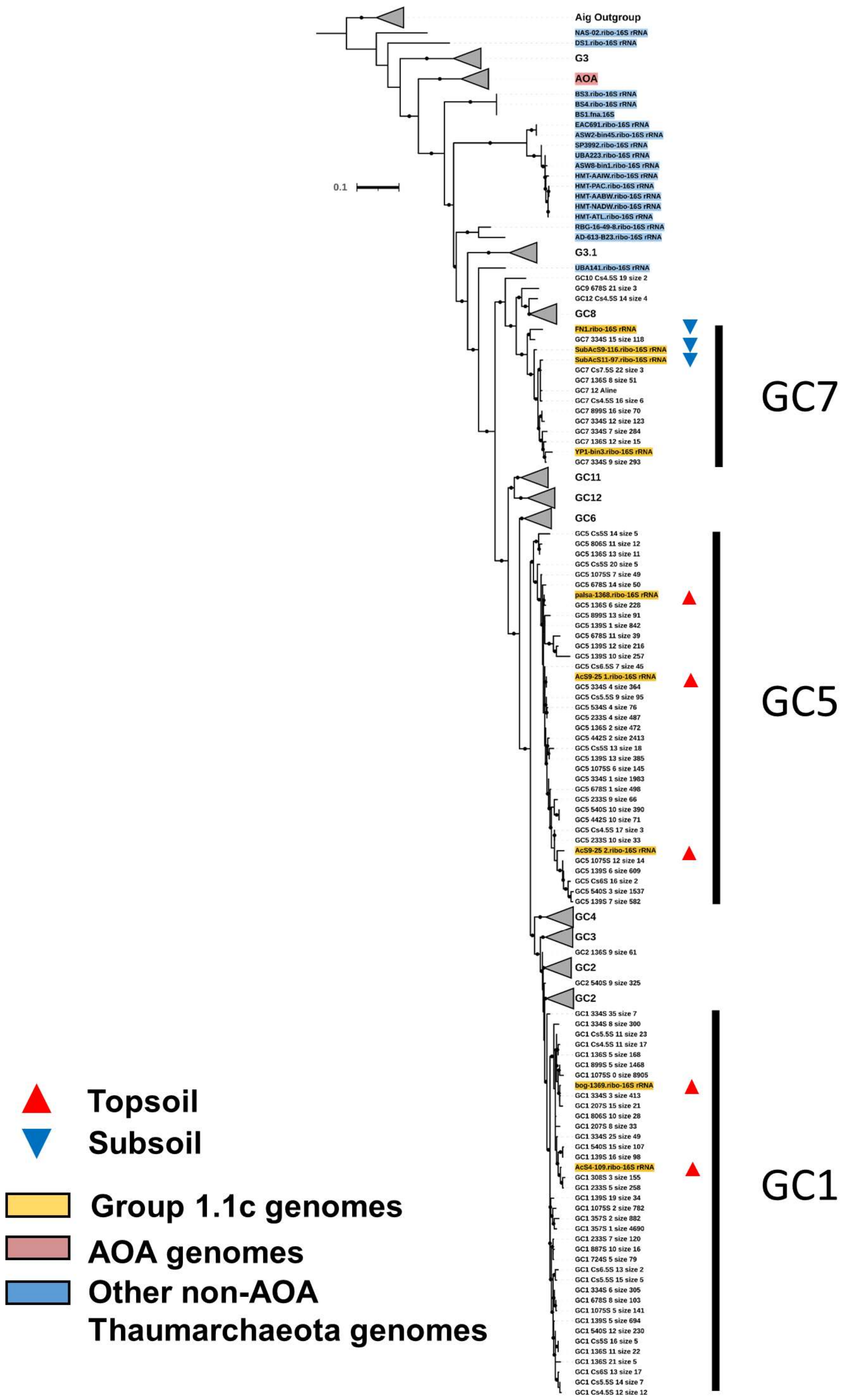


**Supplementary Fig. 2. 16S rRNA gene tree of Group I.1c and related sequences.** This maximum-likelihood tree was created using 16S rRNA genes extracted from the studied genomes and combined with the 16S rRNA database of soil Thaumarchaeota presented in Vico Oton et al 2016[19] (587 column alignment). Dots indicate branches with >70% of 1,000 UFBoot replicates.

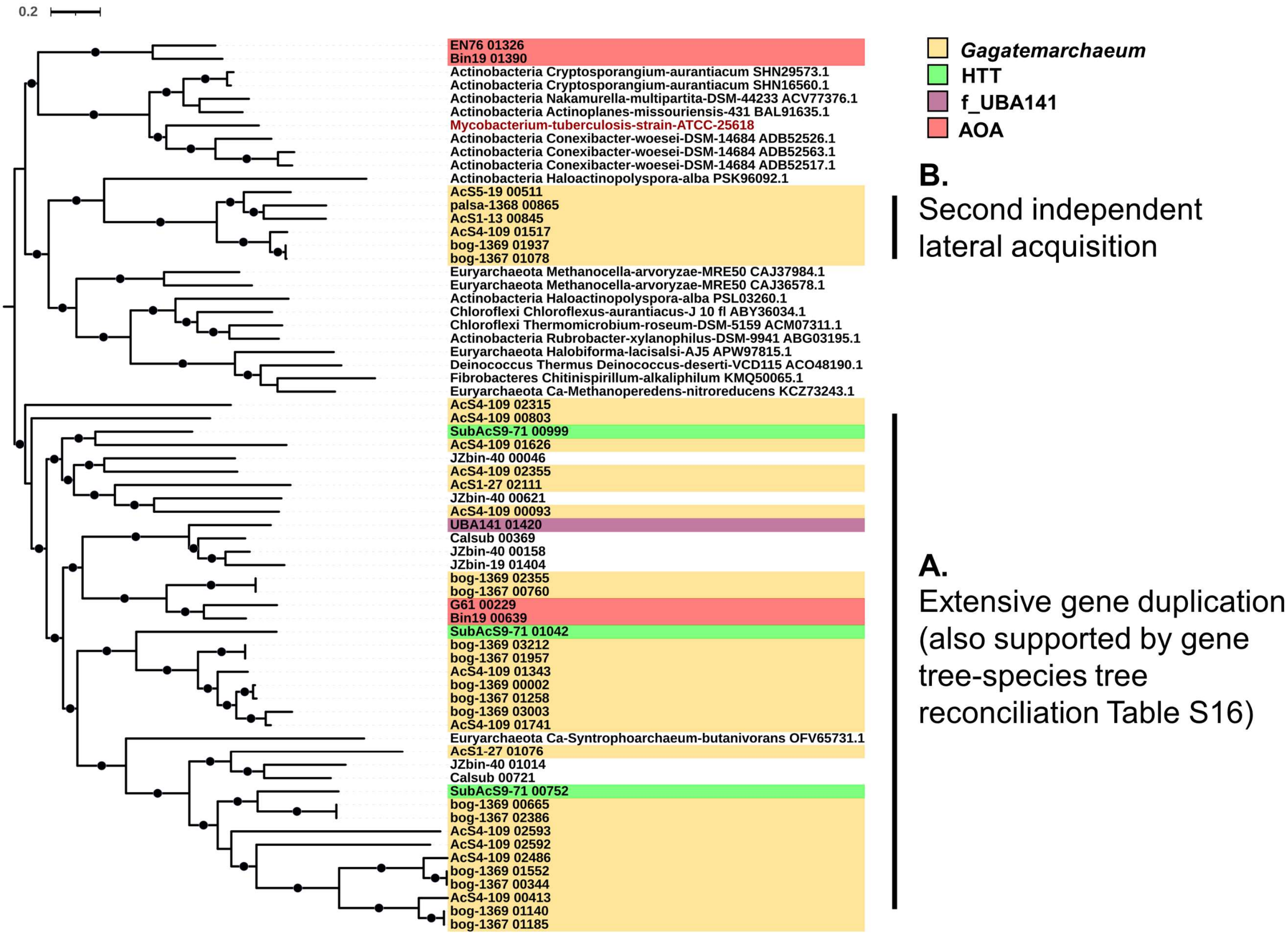


**Supplementary Fig. 3. Phylogeny of F420-dependent glucose-6-phosphate dehydrogenase.** This gene family was present in multiple copies in the topsoil Group I.1c genomes, owing to (A) extensive gene duplication throughout the evolution of *Gagatemarchaeum* and (B) a second independent lateral acquisition. The leaf in red is the experimentally validated gene. Dots indicate branches with >70% of 1,000 UFBoot replicates. The ML tree was created using LG+R4 and rooted with minimal ancestor deviation (MAD).

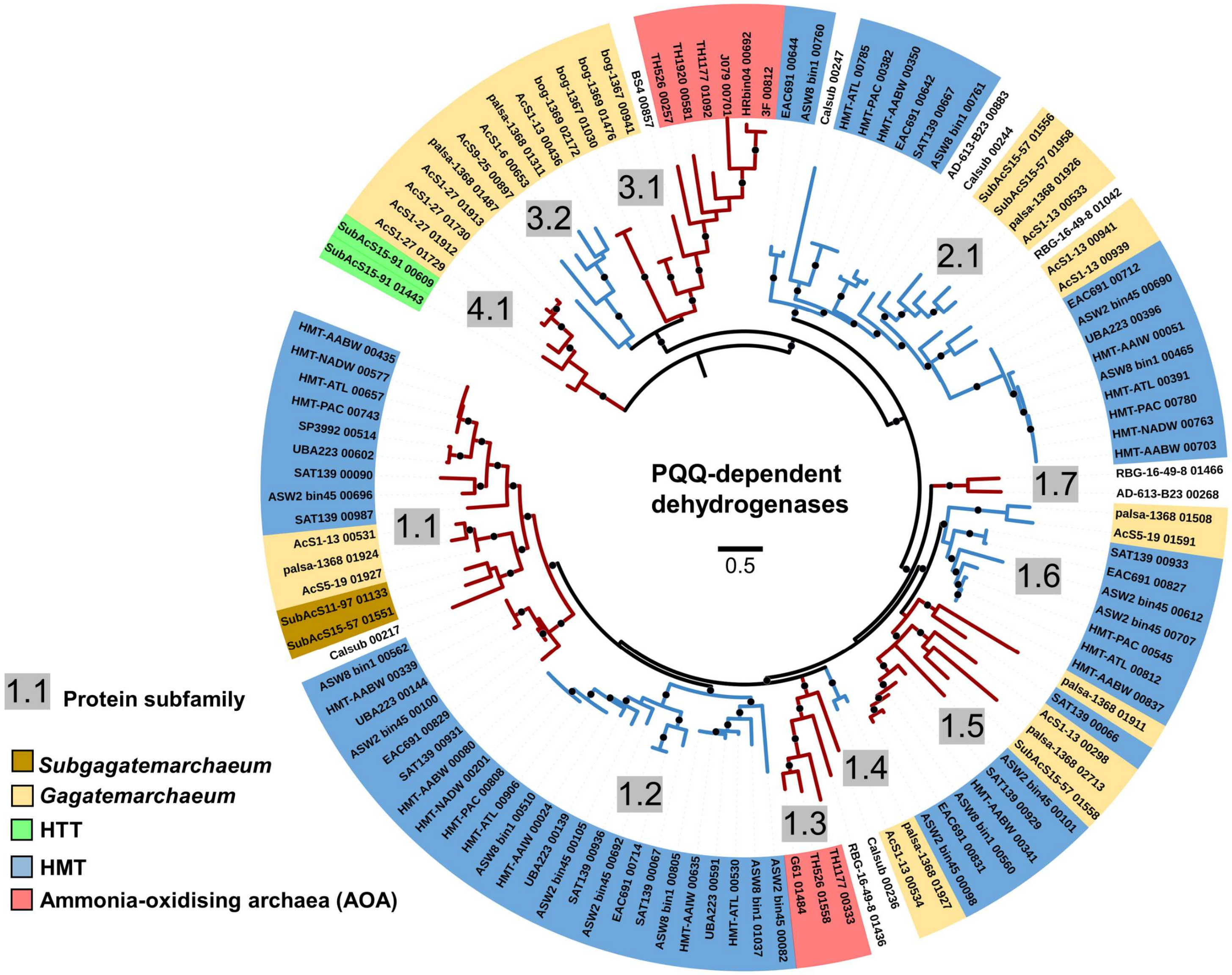


**Supplementary Fig. 4. Phylogeny of Thaumarchaeota PQQ-dependent dehydrogenases.** Dots indicate branches with >70% of 1,000 UFBoot replicates. The alternating blue and red branches indicate different subfamilies as determined by average pairwise distance between leaves. Tree was rooted using minimal ancestor deviation (MAD).

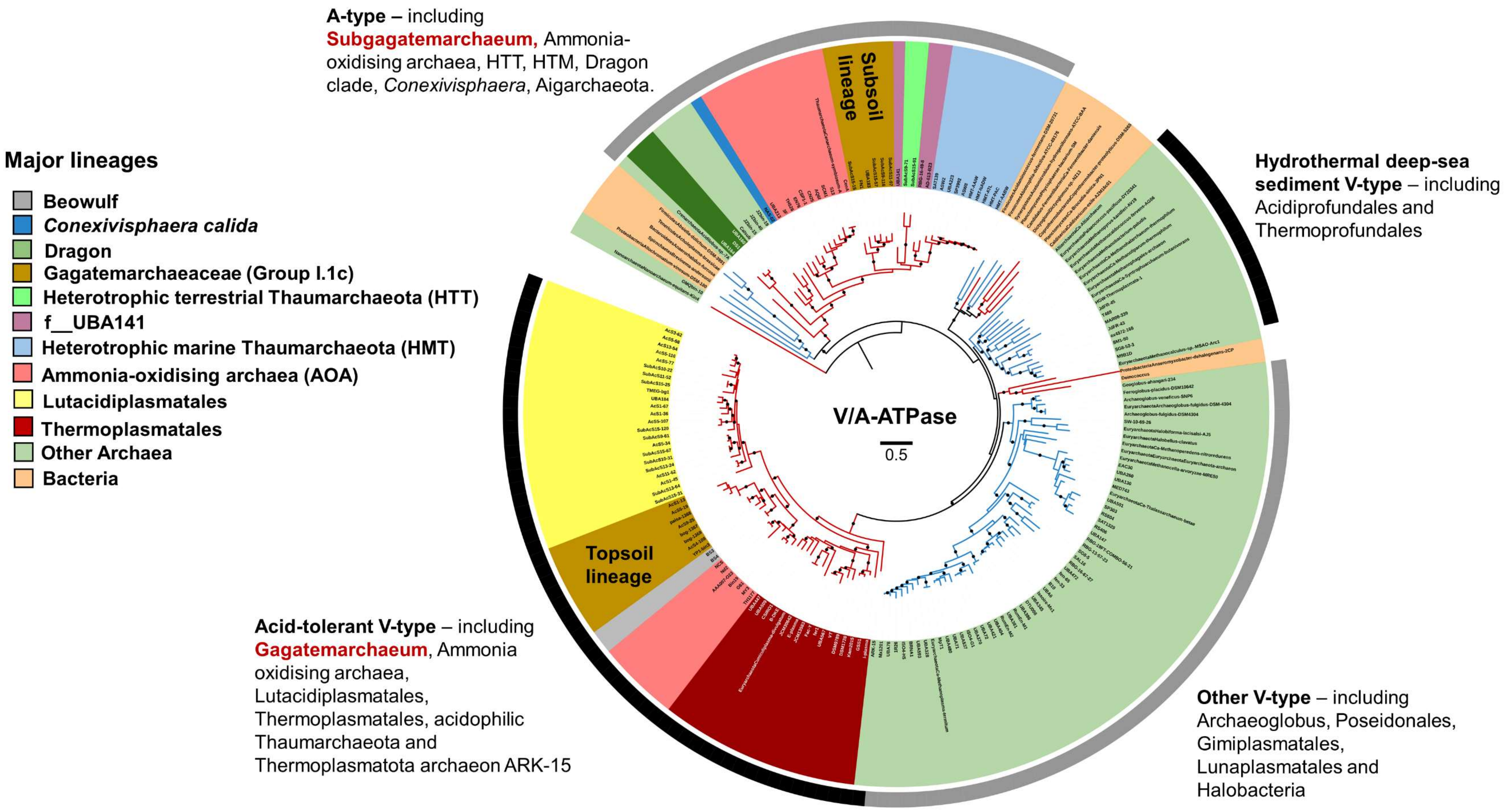


**Supplementary Fig. 5. Phylogeny of the V/A-ATPase.** Sequences from *Gagatemarchaeum* clustered with acid tolerant archaea, whereas those of *Subgagatemarchaeum* clustered with HMT, HTT and non-acidophilic AOA. The three largest subunits of V/A-ATPase (*atpA*, *atpB* and *atpI*) were individually aligned and then concatenated into a single partitioned supermatrix. A supermatrix tree was then estimated using the best fitting model for each partition and rooted using minimal ancestor deviation (MAD). Dots indicate branches with >70% of 1,000 UFBoot replicates. The alternating blue and red branches indicate different subfamilies of V/A-ATPase as determined by average pairwise distance between leaves.

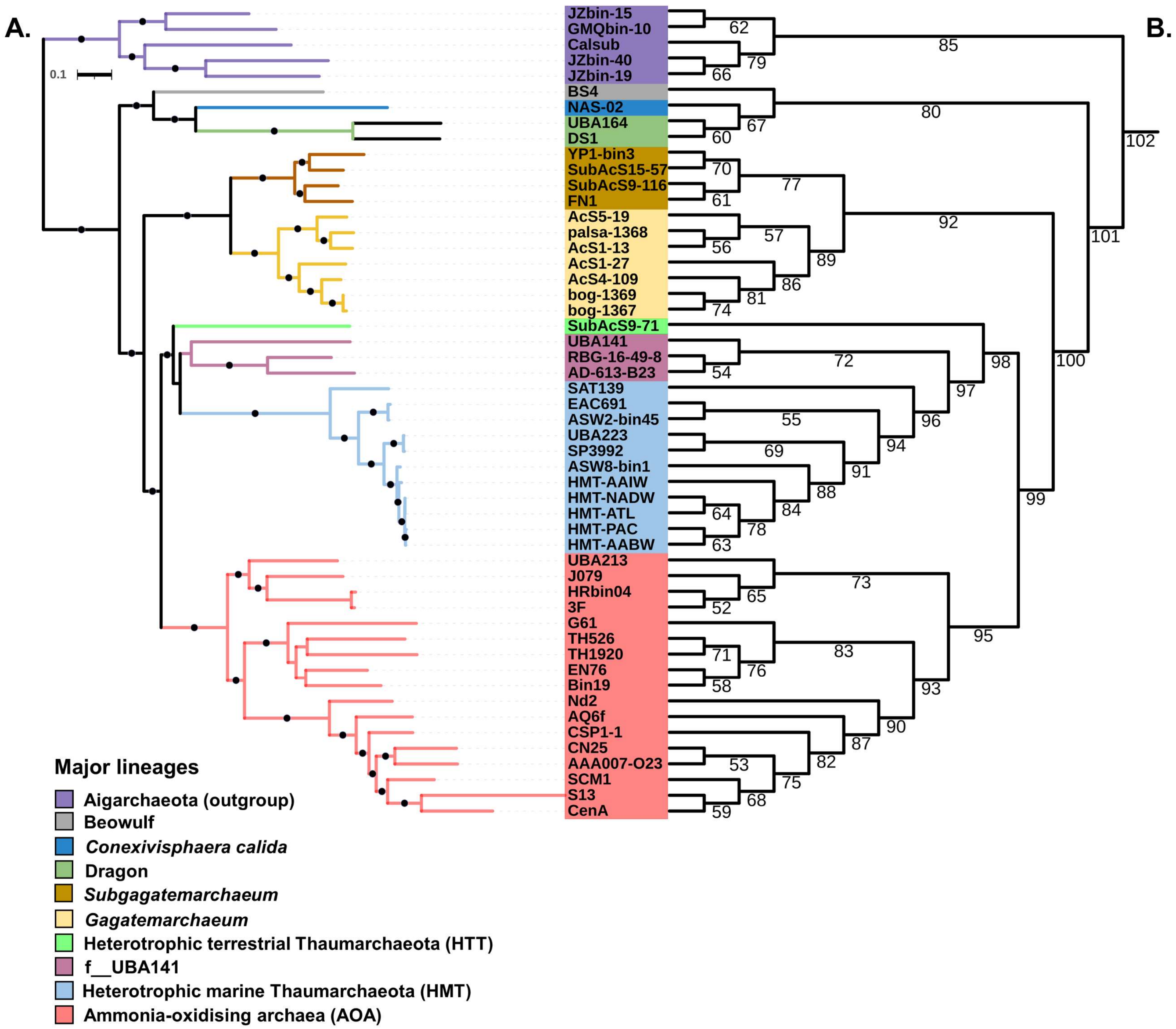


**Supplementary Fig. 6. Phylogeny of higher-quality genomes (A) and corresponding branch numbers (B).** The ML tree was predicted with the LG+C60+F model. Dots indicate branches with >95% of 2,000 UFBoot and 1,000 SH-aLRT replicates**.**

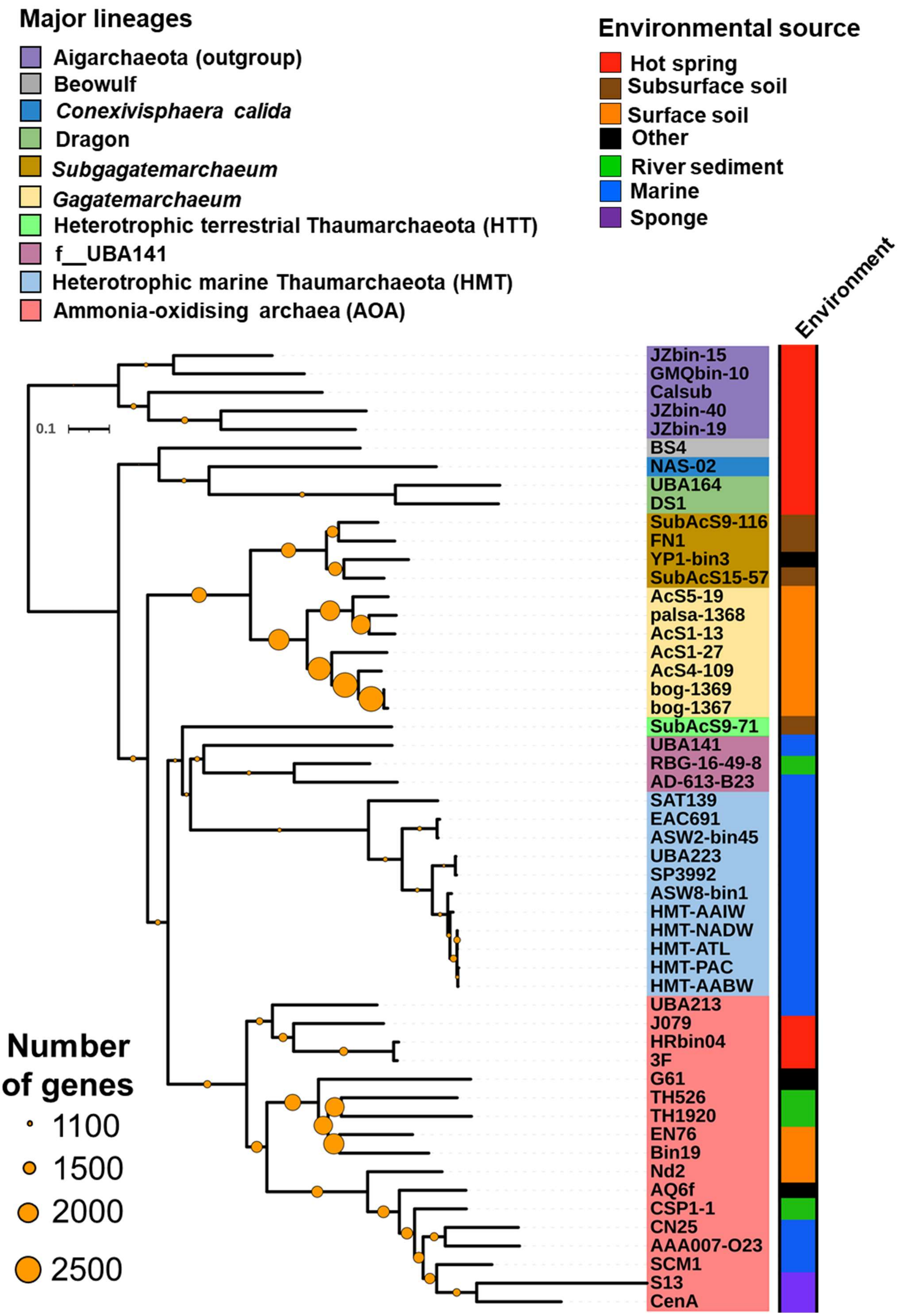
Major lineages
Aigarchaeota (outgroup)
Beowulf
Conexivisphaera calida
Dragon
Subgagatemarchaeum
Gagatemarchaeum
Heterotrophic terrestrial Thaumarchaeota (HTT)
f__UBA141
Heterotrophic marine Thaumarchaeota (HMT)
Ammonia-oxidising archaea (AOA)
Environmental source
Hot spring
Subsurface soil
Surface soil
Other
River sediment
Marine
Sponge
Environment
0.1
JZbin-15
GMQbin-10
Calsub
JZbin-40
JZbin-19
BS4
NAS-02
UBA164
DS1
SubAcS9-116
FN1
YP1-bin3
SubAcS15-57
AcS5-19
palsa-1368
AcS1-13
AcS1-27
AcS4-109
bog-1369
bog-1367
SubAcS9-71
UBA141
RBG-16-49-8
AD-613-B23
SAT139
EAC691
ASW2-bin45
UBA223
SP3992
ASW8-bin1
HMT-AAIW
HMT-NADW
HMT-ATL
HMT-PAC
HMT-AABW
UBA213
J079
HRbin04
3F
G61
TH526
TH1920
EN76
Bin19
Nd2
AQ6f
CSP1-1
CN25
AAA007-O23
SCM1
S13
CenA
Number of genes
1100
1500
2000
2500

**Supplementary Fig. 7. Genome expansion transitioning into terrestrial environments.** Copy number of ancestor gene content reconstructions estimated using gene tree-species tree reconciliation are indicated.

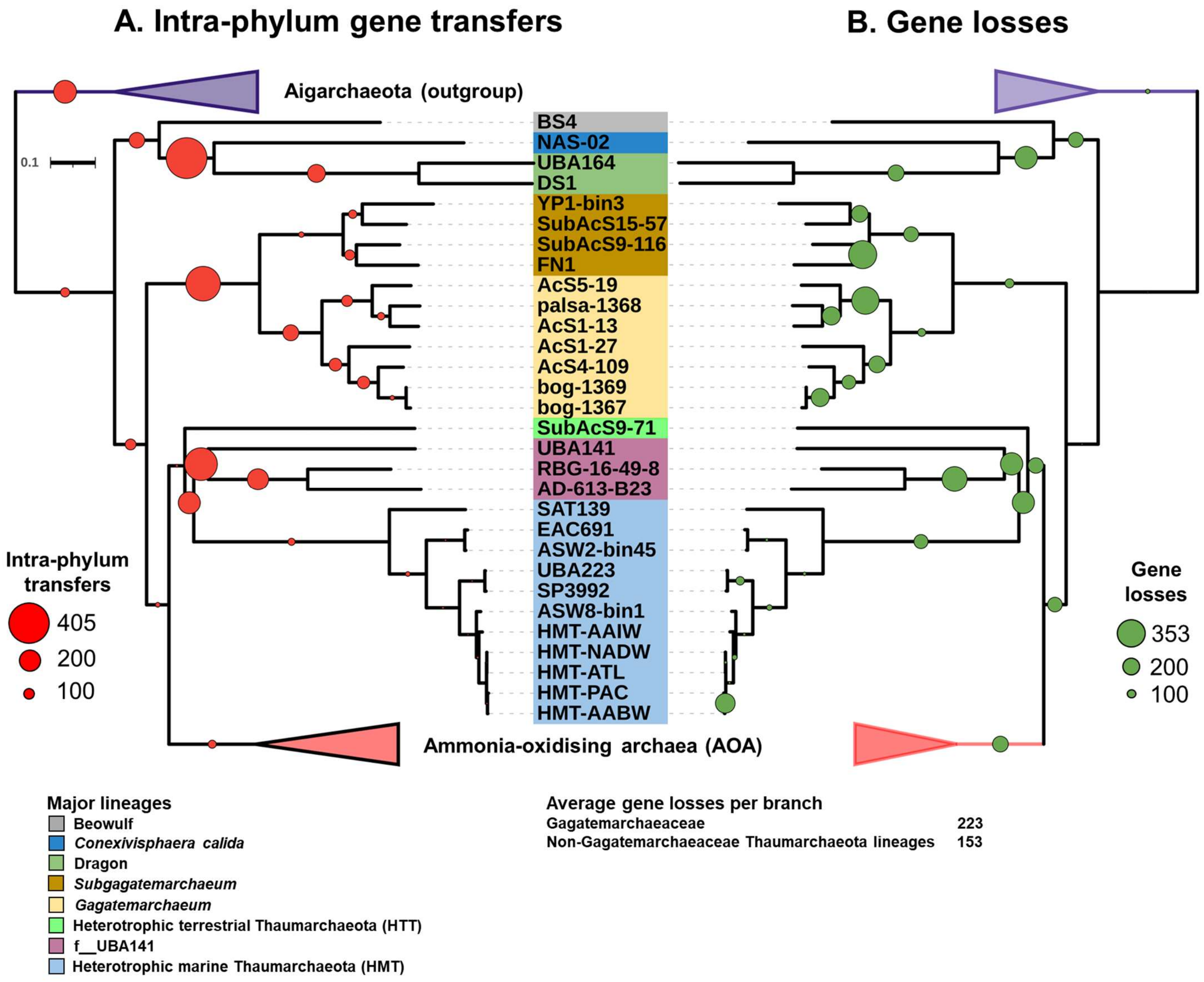


**Supplementary Fig. 8. Acquisition of genes by intra-phyla transfer and gene losses along non-AOA lineages.** The quantitative and qualitative predictions of the genome content changes were estimated across the Thaumarchaeota history using a gene tree-species tree reconciliation approach. Scale numbers indicate the range of the predicted number of events for a given mechanism and circle sizes are proportional to the number of events.

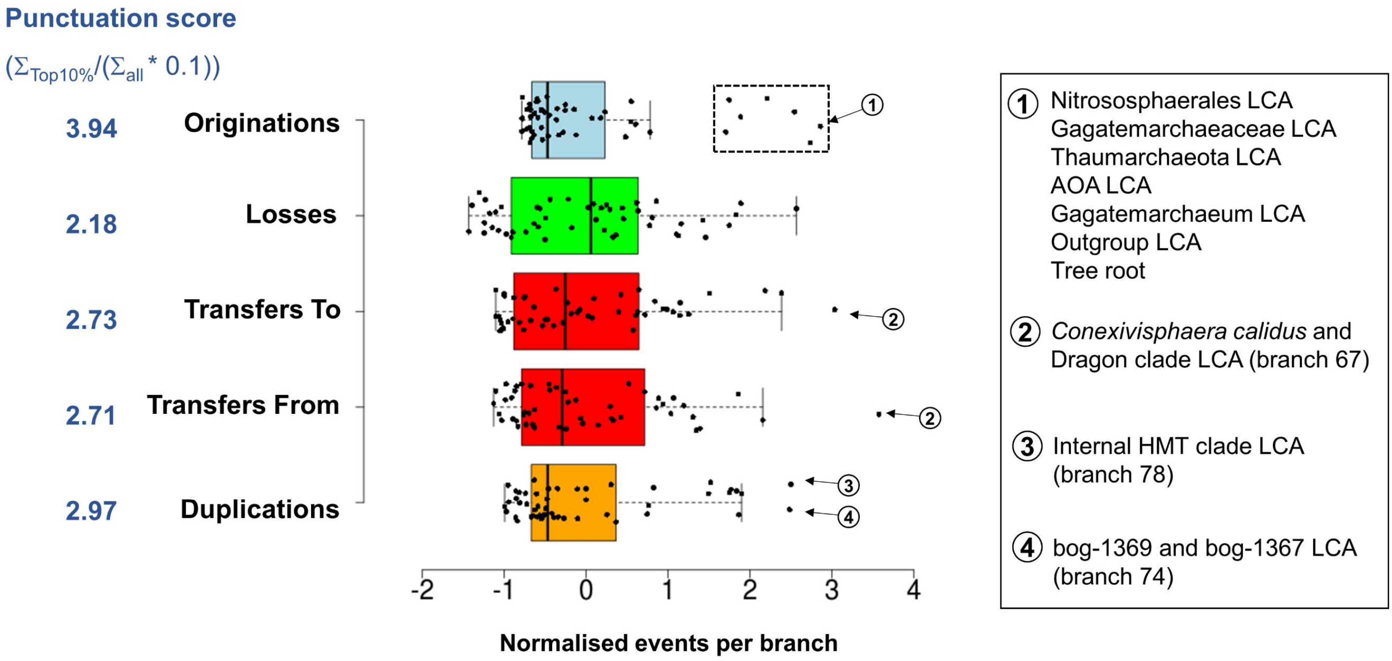


**Supplementary Fig. 9. Distribution of mechanism of gene content change across** Thaumarchaeota evolution. Boxplots represent normalised events per branch ((events per branch - μ)/σ) for each mechanism. Numbered circles mark ancestors (species tree branches) with the highest numbers of events. Horizontal lines within boxes indicate the medians, box boundaries indicate the 1st and 3rd quartiles, whiskers indicate the minima and maxima, and points beyond these whiskers are outliers. A punctuation score is measured for each given mechanism. It represents the sum of events in the 10% of branches with the highest event numbers divided by 10% of the sum of events into all branches ($\Sigma_{\text{events in top10\%}}/(\Sigma_{\text{events in all branches}} * 0.1)$).
